# TCRζ-Driven Pre-Signaling Organization of Lck in Rab11^+^ Endosomes Shapes TCR Activation

**DOI:** 10.1101/2025.09.25.678476

**Authors:** Konstantina Karpouzou, Nikolaos Koutras, Ioannis Tyritidis, Evangelos Tsioupros, Adriana Kotini, Vasileios Roukos, Konstantina Nika

## Abstract

T cell activation relies on the precise spatiotemporal regulation of TCR signaling at the Immunological Synapse, where vesicular trafficking coordinates the delivery of key signaling molecules. While endosomal pools of Lck and its immediate substrate, the TCRζ chain, have been associated with TCR signaling competence, the mechanisms underlying the regulation and synchronization of their trafficking routes remain unresolved.

In this study, we simultaneously traced the endosomal localization dynamics of Lck, ζ chain and the TCR in unperturbed cells and under conditions that preserve the endosomal network integrity. We identified a previously unrecognized plasma membrane resident pool of ζ that exists independently of the TCR complex yet remains competent for phosphorylation and ZAP-70 recruitment. This “standalone” ζ population drives the recruitment of Lck and CD45 into Rab11-positive endosomes, establishing pre-assembled, signaling-primed platforms that could provide the means of sustained signaling without requiring continuous receptor engagement.

Our findings advance the understanding of how vesicular ζ and Lck pools coordinate to shape TCR activation threshold and signaling persistence. Beyond the basic understanding of TCR signaling, these insights hold translational importance for optimizing the design and signaling efficiency of T cell immunotherapies.

## Introduction

T cell activation is triggered by binding of the TCR to cognate peptide-MHC (Major Histocompatibility Complex) molecules displayed by antigen presenting cells (APCs). From the biochemical perspective, initiation of signalling cascades is marked by the phosphorylation of conserved ITAM (Immunoreceptor-based Tyrosine Activation Motifs) tyrosines located in the cytoplasmic tails of the receptor complex, a process tightly regulated by the Src-family kinase (SFKs) Lck. Phosphorylated ITAMs provide anchoring motifs for the recruitment and subsequent activation of ZAP-70, which in turn phosphorylates the adapter protein LAT. Phosphorylated LAT acts as a scaffold for signalosome formation, progressing to signal amplification and diversification and ultimately leading to full T cell activation (1). From the spatial perspective, TCR signaling is initiated, regulated and sustained at the Immunological Synapse (IS), a precisely organized structure formed at the interface between T cells and APCs (2). The orchestrated formation and functionality of the IS depends on time-dependent accumulation, distribution, disassembly and dynamic reorganization, of supramolecular complexes enriched in TCRs, co-receptors and several signaling mediators and adaptors (3). All these events are largely mediated by highly divergent endosomal vesicular transport networks (4). Several key regulators of T cell responses such as Lck, TCRζ, ZAP-70 and LAT are mobilized from vesicular compartments and accumulate at the IS upon TCR engagement (5,6). Moreover, ligand-engaged but also bystander TCRs undergo rapid endocytosis and recycling to the plasma membrane (PM) via endosomal networks, with part of the internalized species being degraded in lysosomes. This behavior also applies to unligated receptors which constitutively follow specialized routes of endocytosis, degradation and recycling. Synchronously, these mechanisms fine-tune TCR surface expression levels, setting the thresholds for signal transduction amplitude, sustainment and termination, and ultimately shape T cell responses (7–11)

The TCR is equipped with a pMHC binding compartment formed by the clonotypic α and β chains and an ITAM-bearing signal transducing unit composed of the invariant CD3γε and CD3δε heterodimers, and a ζζ homodimer. Assembly of the TCR is initiated in the Endoplasmic Reticulum and completed in the Golgi apparatus by the association of the ζ homodimer. The ζ chain constitutes the limiting factor for both TCR transport to the cell surface but also for stable expression of the complex at the PM (12).

Although TCR and CD3 are often concurrently downregulated, the intracellular trafficking and fate of the ζ subunit is, at least in part, uncoupled from the rest of the complex, characterized by prolonged and more readily detectable residency within endosomes. Furthermore, dissociation of ζ from the rest of the TCR, alongside a continuous exchange with TCR-CD3 partial complexes has been reported (13–16). Endosomal ζ has been the focus of extensive investigation, with numerous studies—occasionally reporting divergent results—implicating Rab4, Rab5, Rab8, Rab11, VAMP3, VAMP7, and IRAP, as its colocalization partners (4,6,17–19). Vesicular ζ, has been shown to originate from internalized cell surface-resident ζ molecules, to dynamically exchange with the PM and to provide a sustained supply of ζ molecules to the Immunological synapse. Notably, endosomal ζ -particularly the fraction colocalizing within Rab11 has been found to be tyrosine phosphorylated in resting and stimulated cells, an event attributed to the co-residence of Lck within the same compartment (17,19). Accordingly, the biological implications of Lck residency in the Rab11-decorated endosomal compartment (Rab11^+^ EC) have been extensively characterized (6,20,21). Collectively, chemical treatments or genetic deletion of key recycling endosomal regulators have shown that disrupting the vesicular trafficking of Lck and ζ impairs their recruitment to the immunological synapse and compromises TCR signalling (17,19–21). However, these approaches have largely relied on fluorescently tagged proteins or synthetic ζ chain reporters, without fully addressing how such modifications may affect their incorporation into the native TCR complex. Moreover, chemical or genetic perturbations of endosomal regulators may introduce unintended alterations in endosomal network architecture and function. Lastly, despite extensive evidence highlighting the biological importance of the vesicular Lck and ζ pools, the extent to which their trafficking dynamics are interconnected and co-regulated remains incompletely understood.

In this work, we concurrently trace the endosomal localization dynamics of endogenous TCR, Lck and ζ chain in unperturbed cells, under conditions that preserve the integrity of the endosomal network. Our findings reveal a previously unrecognized role of TCRζ as a driver for Lck and CD45 residency within the Rab11^+^ endosomal compartment (EC). We provide evidence that this property is accredited to a PM-resident ζ pool that is physically uncoupled from the TCR complex yet retains signalling competence. These insights contribute to advancing our understanding on TCR sensitivity and rapid responses and hold particular relevance within the context of chimeric receptor-based cell therapies

## Results

### Exclusion of Lck from Rab11^+^ endosomes correlates with reduced TCR signalling output

To gain insight into the mechanisms governing the endosomal trafficking of the most proximal TCR signalling mediators, we initiated our investigation by examining the contribution of the Lck SH4 domain, a 10–amino acid long N-terminal segment. SH4 domains enable SFK anchoring to membranous structures; they invariably undergo co-translational myristoylation and post-translational palmitoylation -albeit with different patterns- and can contain basic amino acids that facilitate and stabilize SFK interactions with membranes. These unique biochemical features confer distinct subcellular localizations and spatial membrane distributions to SFKs ultimately regulating their function (22–26). Our study incorporated chimeric constructs in which LckSH4 was substituted by the corresponding sequences of Fyn and Lyn (designated Fyn-Lck and Lyn-Lck) alongside the SH4 deletion mutant (LckΔSH4) serving as negative control for LckSH4 function (Fig. 1A). Wild-type Lck (LckWT) and chimeras were inducibly expressed in the Lck-devoid jurkat variant JCaM1.6 under the control of Doxycycline (Dox) (FigS1A).

**Figure 1.**
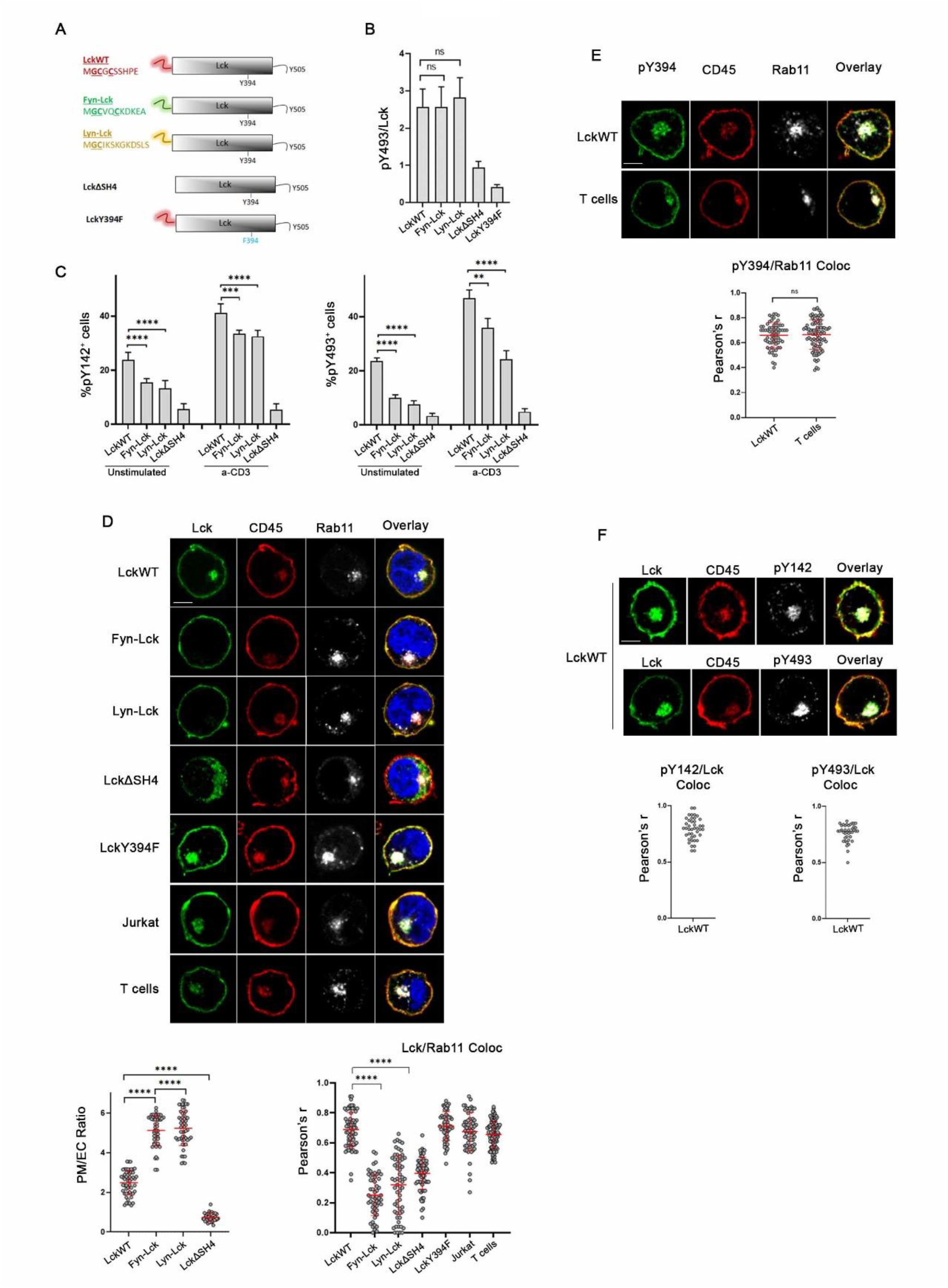
Characterization of Lck and ζ chain localization within Rab11^+^endosomes. A. Schematic illustration of LckWT, Fyn-Lck, Lyn-Lck, and the ΔSH4 deletion and Y394F mutants. The SH4 domain amino acid sequences are listed, myristoylation (Glycine) and palmitoylation (Cysteine) sites are in bold and underlined. B. LckSH4 substitution does not compromise the levels of constitutively active Lck. JCaM1.6 cells expressing the indicated LckWT, chimeric and mutant forms were stained for total Lck expression and for its active form (pY394-Lck) and analysed by FACS (Gating strategy described in Fig.S1A). MFI values for each antibody were used to calculate absolute levels of active Lck pY394/Lck ratio) for each construct. The graph displays pooled data from four independent experiments C. LckSH4 substitutions reduce the TCR proximal signaling output. JCaM1.6 cells expressing the indicated LckWT, chimeric and mutant forms were either left unstimulated or incubated with 1μg/ml a-CD3 for 2min at 37×C. Cells were stained with a-Lck and antibodies against the phosphorylated forms of ζ chain ITAM Y142 or the activating Y493 of ZAP-70 and analysed by FACS. Graphs show the frequency of cells displaying phosphoantibody fluorescence levels exceeding the -Dox baseline, within Lck^+^-gated populations (gating strategy described in Fig.S1B). Data from four independent experiments. D. Confocal microscopy of JCaM1.6 cells expressing the indicated constructs, jurkat cells and human CD4^+^ T cells, stained for Lck (*green*) and Rab11 (grey). PM and nucleus are defined by CD45 (*red*) and DAPI staining (*blue*) respectively. *Left graph*: PM/EC ratio of a-Lck fluorescence intensity. ROIs for each compartment were defined by CD45 and Rab11 staining respectively. *Right* graph: Collective colocalization analysis results for a-Lck and a-Rab11 staining as indicated. n ≥ 40 cells from two independent experiments E. Constitutively active Lck can be detected in the EC. Human CD4^+^ T cells and JCAM1.6 cells expressing LckWT were stained for active Lck (pY394-green) CD45 (red) and Rab11 (grey). *Bottom graph*: colocalization scores for pY394 and Rab11 staining. n≥60 cells from three independent experiments. F. Endosomal Lck colocalizes with activated ζ and ZAP70. JCAM1.6 cells expressing LckWT were stained for Lck (green), CD45 (red) and *Top panels*: pY142 (grey), or *bottom panels:* pY493 (grey). *Bottom graphs*: colocalization scores for a-Lck and the indicated phosphoantibodies. n ≥ 40 cells from three independent experiments. Scale bars, 5μm. Unpaired Student t test; mean +/- standard deviation [SD]; ns: not significant, **P < 0.01, ****P < 0.0001.

Substitution of the SH4 domain did not compromise the levels of active Lck (pY394-Lck) (Fig.1B and Fig.S1A) whereas, as previously shown, the SH4 deletion mutant (LckΔSH4) and LckY394F (Fig.1B and S1B), where the activating tyrosine is replaced by phenylalanine were largely devoid of kinase activity (24). Unexpectedly, despite equivalent levels of pY394-Lck, the chimeric forms were significantly less effective in driving ζ chain ITAM Y142 (pY142-ζ) and ZAP-70 Y493 (pY493-ZAP) phosphorylation in a-CD3 (clone UCHT1) stimulated cells (Fig. 1C and Fig.S1C).

This pattern appeared more pronounced in unstimulated samples, where several groups, including ours, have demonstrated that Lck overexpression or deregulated activity can autonomously initiate signalling cascades independently of receptor ligation (27–29). These data unveil an additional role of SFK SH4 domains in optimal substrate juxtapositioning.

To exclude the possibility that the signaling inefficiency of Fyn-Lck and Lyn-Lck was due to their impaired localization at the plasma membrane (PM) -the compartment where the primary Lck substrates (TCR ITAMs) reside- we performed confocal microscopy analyses. As previously shown (20,21), Lck was distributed between the inner leaflet of the plasma membrane, and the Rab11^+^ EC, whereas LckΔSH4 was predominantly cytoplasmic (Fig.1D). On the contrary, Fyn-Lck and Lyn-Lck were undetectable in the intracellular space, displaying a significantly higher PM/EC ratio than LckWT, indicative of their exclusive PM residence. Accordingly, the SH4 chimeras did not colocalize with Rab11(Fig.1D). Interestingly the Rab11 colocalization scores were indistinguishable between LckWT and the kinase-inactive mutant LckY394F, stating that Lck kinase activity is not a prerequisite for endosomal partitioning.

To delineate the potential association of endosomal Lck with signaling outputs, we proceeded to the characterization of Lck and its co-resident molecules within Rab11^+^ structures. Confocal microscopy analyses, in agreement with previous reports (6,19,20), confirmed that the endosomal Lck pool harbored its constitutively active form (pY394-Lck) in LckWT-expressing cells and human CD4 T cells (Fig.1E). Endosomal Lck also strongly colocalized with pY142-ζ and pY493ZAP70 (Fig.1F) in unstimulated cells overexpressing LckWT.

### TCRζ expression is required for Lck trafficking to the Rab11^+^ EC

The presence of internalized phosphorylated ζ chain (pY142-ζ) is consistent with the concept of SFK activity-dependent TCR endocytosis (14,30). However, strong Rab11-ζ colocalization scores were obtained in parental JCaM1.6 cells and in jurkat cells incubated with the pan-SFK inhibitor PP2 for 18h (Fig.S2A). Furthermore, ζ/Rab11 colocalization was not compromised in cells expressing Fyn-Lck and Lyn-Lck (Fig.2A,2C and S2B). These data demonstrate that the expression of Lck per se, its endosomal presence or SFK enzymatic activity (i.e. ITAM phosphorylation) are not prerequisites for ζ chain residence in the Rab11^+^ EC. TCRζ also strongly colocalized with Rab11 in the J76 jurkat subline that is devoid of endogenous TCRα and β chains (Fig.2A, 2C and Fig.S2A), suggestive of Rab11^+^ endosomal residency being an inherent ζ chain trait that can occur independently of receptor internalization. In contrast to ζ, CD3ε did not colocalize with Rab11 neither in human T cells nor in the LckWT line (Fig,2B, 2C), corroborating the concept of divergent trafficking amongst the TCR-CD3 subunits and the ζ chain.

**Figure 2.**
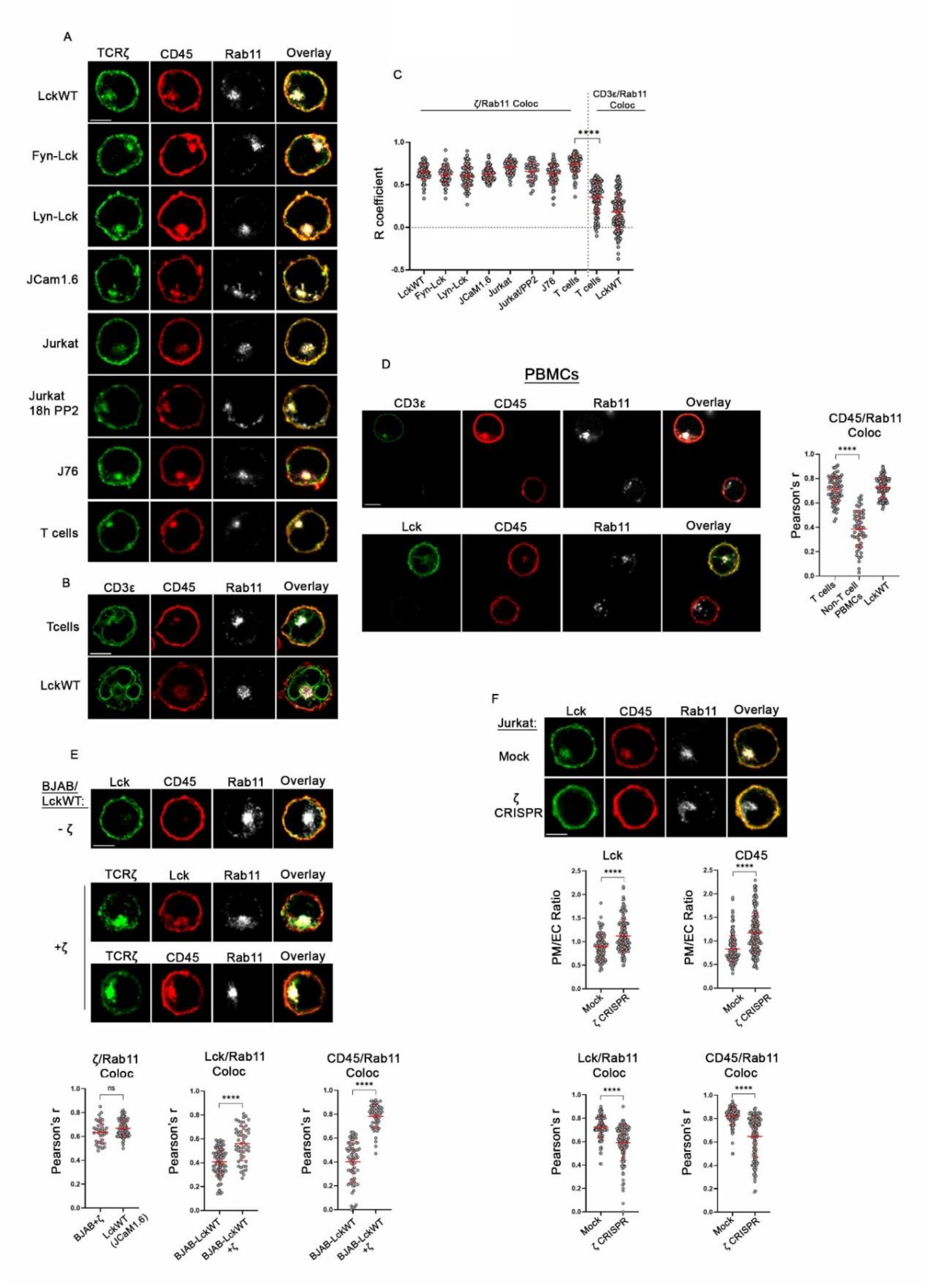
TCRζ drives Lck endosomal residency. A. A fraction of ζ chain molecules is localized in Rab11^+^ endosomes. Confocal microscopy analysis of JCaM1.6 lines expressing LckWT, Fyn-Lck, Lyn-Lck, JCaM1.6 cells, Jurkat cells untreated or cultured in the presence of the pan-SFK inhibitor PP2 for 18h, the surface TCR-devoid J76 subline and human CD4 T cells stained for TCRζ (green), CD45 (red) and Rab11 (grey). n≥40 cells from three independent experiments. B. CD3ε does not co-reside with ζ in Rab1^+^ EC. Human CD4 T and JcaM16-LckWT cells were stained for CD3ε (green), CD45 (red) and Rab11 (grey). n ≥100 cells from three independent experiments. C. Graph of combined colocalization data between Rab11 and TCRζ or CD3ε from A and B, as indicated. D. Endosomal CD45 is absent from non-T cell PBMCs. Freshly isolated PBMCs were stained with CD45 (red) and Rab11 (grey), along with either CD3ε, or Lck (green) as indicated, to distinguish T cells from other lymphocyte populations. Adjacent graph shows collective colocalization analysis scores for CD45 and Rab11 in T cells and non-T cell PBMCs within the same slide and in the LckWT line (images displayed in Fig.1D, 1E and Fig.2A). n ≥50 cells from three independent experiments. F. Ectopic ζ expression in BJAB promotes Lck and CD45 endosomal residency. *Top panel*: BJAB-LckWT cells stained for Lck (green), CD45 (red) and Rab11 (grey). *Bottom panels*: BJAB-LckWT cells co-expressing the TCRζ chain were stained for ζ (green), Rab11 (grey) and either Lck or CD45 (red) as indicated. Low magnification images shown in Fig.S2D. *Bottom graphs*: Colocalization analyses scores as indicated F. ζ chain silencing diminishes the endosomal Lck and CD45 pools. Confocal images of CRISPR- or mock-treated jurkat cells stained for Lck (green), CD45 (red) and Rab11 (grey). Low magnification images shown in Fig.S2F. Graphs depict PM/EC ratio for Lck and CD45 (top panels) or their colocalization with Rab11 (bottom panels) as indicated. n ≥100 cells from three independent experiments. Scale bars, 5µm. Unpaired Student t test; mean +/- SD; ****P < 0.0001, ns, not significant

Throughout all imaging analyses, CD45 staining was employed to delineate the plasma membrane boundary. Surprisingly, we consistently observed that CD45 exhibited a pronounced vesicular presence (Fig. 1D-F, Fig.2A, 2B). Endosomal CD45 was not the result of antibody cross-reactivity, as its presence was consistent across various staining combinations, but more importantly because its colocalization with Rab11 could not be detected in non-T cell PBMCs (Fig. 2D). Intrigued by the scenario that a Rab11-centered signaling module might be a unique feature of T cells and directly related to inherent properties of the TCRζ chain, we transitioned to the B cell line BJAB, which is devoid of the TCR apparatus. We reasoned that if Lck and/or CD45 positioning within the Rab11^+^ compartment was a T cell signature event, mediated by the process of ζ chain internalization, this phenomenon would not withstand in the B cell environment. For these experiments we used BJAB cells expressing LckWT under the control of Dox. In a previous study (31) we fully characterized ectopic Lck in this B cell environment showing that the kinase was constitutively active, similarly to its endogenous counterpart in T cells, and biologically functional as it could encounter and phosphorylate BCR ITAMs and trigger downstream signaling cascades. Nevertheless, both Lck and CD45 were completely absent from Rab11^+^ structures in BJAB cells (Fig.2E and Fig.S2D). This pattern was overturned upon transduction with human TCRζ chain. Ectopically expressed ζ strongly colocalized with Rab11 regardless of the presence of absence of Lck. Moreover, ζ expression was sufficient to drive the redistribution of Lck and CD45 to the Rab11^⁺^ endosomal compartment (Fig.2E and Fig.S2C, S2D).

These data were further corroborated by CRISPR/Cas9-mediated silencing of endogenous TCRζ in Jurkat cells. ζ downregulation was accompanied, as anticipated, by a dramatic drop in levels of surface TCR alongside a significant increase in the PM/EC ratio of Lck and CD45 residency and their severely reduced colocalization with Rab11, compared to the mock-treated controls (Fig.2F and Fig.S2E,S2F)

Overall, these data portray a fraction of ζ chain molecules, uncoupled from the TCR complex, that act as drivers for Lck EC residency. Nonetheless, the origin and trafficking dynamics of Lck and ζ chain endosomal pools in unperturbed cells remains an open question.

### A constitutively TCR-uncoupled ζ pool drives Lck trafficking from the plasma membrane to Rab11^+^endosomes

Based on our data, Fyn-Lck and Lyn-Lck serve as excellent models of exclusively PM-localized Lck, and were used to trace, directly the pool of Lck, and indirectly, of the ζ chain destined for Rab11^+^ endosomal residency. We examined the possibility that despite their endosomal absence in resting cells, Fyn- and Lyn-Lck could undergo internalization upon triggering of the signaling apparatus.

Cells were initially treated with Pervanadate, aiming for a generalized and very robust form of cell activation. By 10min of Pervanadate treatment Fyn-Lck, Lyn-Lck and pY142-ζ could be clearly detected within the Rab11^+^ structures (Fig.3A and Fig.S3A). This was a time-dependent process, weak detection of Lck chimeras to the EC required at least 2-3min of Pervanadate treatment (data not shown). pY142-ζ could not be detected in untreated cells, concordant with very low levels of global steady-state pY142-ζ shown in Fig.1C. In agreement with previous studies (30), pervanadate stimulation was not followed by TCR internalization (Fig.S3D), indicating cohesion of the sTCR complex (15). CD3ε remained well segregated from Rab11 in pervanadate stimulated samples (Fig.S3D).

**Figure 3.**
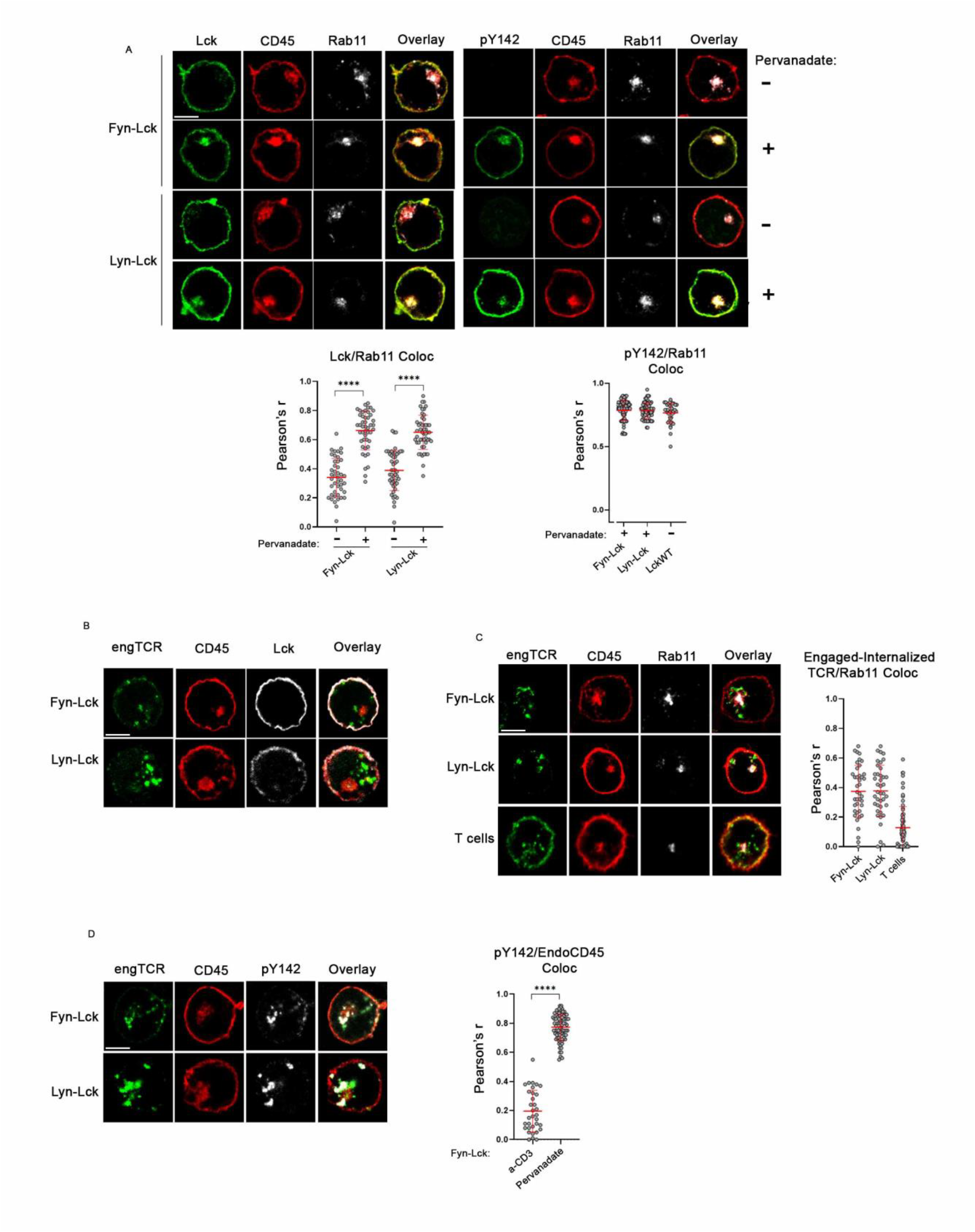
A non-TCR-associated ζ pool is co-trafficking with Lck to the Rab11^+^EC. A. Pervanadate promotes Fyn-Lck and Lyn-Lck translocation, and pY142 presence at the Rab11^+^EC. Cells were either left untreated or incubated with 100μM Pervanadate for 10min and stained with CD45 (red), Rab11 (grey) and either Lck (green) or pY142 (green), as indicted. n ≥40 cells from three independent experiments. *Bottom graphs*: colocalization scores as indicated. In the pY142/Rab11 colocalization graph (right), an additional column was included for comparison purposes, showing Pearson’s r values from LckWT-expressing cells, (originally presented in Fig.1F). B. TCR engagement does not induce Lck translocation to the Ra11^+^EC. The Fyn-Lck and Lyn-Lck lines were or stimulated with 1 µg/μl anti-CD3 for 10min and directly stained with a-mouse 2^ary^ antibody (green) to exclusively visualize the engaged TCRs, CD45 (red) and Lck (grey). Representative images from three independent experiments. C. Internalized/engaged TCRs do not transition to the Rab11^+^EC. Fyn-, Lyn-Lck expressing cells or human CD4 T cells, were stimulated as in (B) and stained with a-mouse 2^ary^ antibody (green), CD45 (red) and Rab11 (grey). *Adjacent graph*: colocalization scores between engaged/internalized TCRs and Rab11. n ≥40 cells from two independent experiments D. TCR engagement does not induce endosomal ζ phosphorylation. Fyn-, Lyn-Lck expressing cells were stimulated as in (B) and stained with a-mouse 2^ary^ antibody (green), CD45 (red) and pY142 (grey). n≥30 cells from three independent experiments. *Adjacent graph*: colocalization scores in the Fyn-Lck line, between pY142 and endosomal CD45 (EndoCD45). Due to limitations imposed by antibody species incompatibility, endosomal CD45 was used to indirectly define ROIs for the Rab11^+^EC. For comparison purposes, an additional column was included, showing Pearson’s r values from pY142/CD45 colocalization of unstimulated LckWT cells (Images used for analysis are presented in Fig.1F). Scale bars, 5 µm. Unpaired Student t test; mean +/- SD; ****P < 0.0001

To assess whether the pervanadate-induced endosomal residency of the chimeras was merely a spurious effect of enforced, non-physiological cellular processes triggered by pervanadate, we performed identical experiments in BJAB cells, where in the absence of ectopic ζ expression, Lck is exclusively localized at PM (Fig.2E). Unlike data obtained in the JCaM1.6 lines, pervanadate did not promote Lck internalization in BJAB, with the kinase remaining at the PM (Fig.S3B, S3C). Thus, the pervanadate-induced Rab11^+^EC accumulation of Fyn-Lck and Lyn-Lck in pervanadate-treated cells is ζ chain dependent.

Collectively, these data attest that Rab11^+^ resident Lck originates from the PM, where it has to encounter the ζ chain in order to get co-transported to the Rab11^+^ EC.

It is universally accepted that in TCR-sufficient cells, PM-resident ζ species are embedded in the fully assembled receptor complexes (32,33). Since Rab11^+^ endosomal ζ does not colocalize with CD3ε, it is reasonable to deduce that PM resident Lck co-internalizes with a fraction of ζ molecules that disengage from the receptor complex. We investigated this hypothesis by stimulating Fyn-Lck and Lyn-Lck expressing cells for 10min with the mouse anti-CD3ε antibody, clone UCHT1. These stimulatory conditions have previously been shown to induce significant ζ chain dissociation and consequent surface TCR (sTCR) downregulation (15). anti-CD3 treated samples were fixed, permeabilized and directly incubated with fluorescently conjugated anti-mouse secondary antibody to exclusively trace the fraction of engaged receptors. Coverslips co-stained for Lck or Rab11 revealed that, neither the Lck SH4 chimeras nor the internalized engaged TCRs localized within the Rab11^+^ EC after a-CD3 stimulation (Fig.3B, 3C). Due to species incompatibility restrictions of commercially available antibodies, the Rab11^+^EC in Fig.3B and 3D was indirectly demarcated by endosomal CD45

Complementary staining with anti-pY142 (rabbit) revealed no Rab11^+^ EC enrichment of phosphorylated ζ species in anti-CD3 stimulated cells, although phosphorylated ζ could clearly be detected at the PM and intracellularly colocalized with a proportion of the engaged, receptors (Fig.3D and Fig.S3F). Image analyses showed that 35-40% of intracellular anti-CD3-bound TCRs did not colocalize with pY142-ζ (Fig.S3F), attesting its disengagement from the complex, in agreement with previous estimations (15). This percentage was retained in samples that were co-incubated with pervanadate during the last 2min of a-CD3 treatment -to augment anti-pY142 reactivity-thus excluding overestimations of ζ disengagement due to the presence of unphosphorylated ζ molecules attached to the internalized complexes (Fig.S3E and F).

Collectively these data attest that the fraction of ζ molecules that dissociate from the complex under conditions of TCR engagement, or get internalized together with the ligated receptors constitute distinct pools from the ζ chain fraction that co-transits with Lck to the Rab11^+^ EC.

## Discussion

In this work we revisited the long-lasting question of the mechanisms governing the internalization and recycling dynamics of the proximal TCR signaling machinery.

Endosomal trafficking of the TCR complex, ζ chain and Lck has primarily been studied in the context of post-receptor engagement events, where it has been associated with optimal TCR responsiveness and performance as well as IS functionality. In accordance with these studies, we show that Lck chimeric forms devoid of endosomal residence were less efficient than the WT protein in triggering TCR signaling events. This phenotype was irrespective of ζ chain residency in the EC, highlighting the biological significance of Rab11^+^EC resident Lck.

We provide evidence that in unstimulated T cells, a distinct pool of the TCRζ chain that is spatially separated from the TCR-CD3 subunits, constitutes the driving force for the assembly of a group of signaling mediators within Rab11^+^ endosomal structures. Under conditions of Lck overexpression or pervanadate treatment the Rab11-decorated compartment was enriched in pY142-ζ and pY493-ZAP70, the presence of which was absolutely dependent on the residence of Lck within the same location. Rather surprisingly CD45 also had a strong presence within the Rab11^+^ endosomal structures. Although the trafficking and redistribution of CD45 in distinct compartments of the IS has been extensively investigated (34), imaging analyses have primarily focused on its lateral mobility in planar views of TCR-pMHC interfaces. Indeed, we could only find a single report documenting -via immunofluorescence microscopy- the presence of an intracellular CD45 pool in resting Jurkat cells, suspected to participate in the phosphatase’s turnover at the PM (35). However, this observation was not further pursued and to our knowledge, endosomal CD45 has not been previously reported. In light of our data, a key question emerges: how are the opposing actions of the aforementioned endosomal co-residents coordinated within the context of T cell activation? Studies on the competing phosphorylation-dephosphorylation dynamics of the proximal TCR signaling machinery on lipid bilayers (36) showed that the ζ chain phosphorylation output was dependent on the local concentrations of Lck and CD45. Notably, at equal Lck-CD45 densities, ζ phosphorylation was readily detectable and was further enhanced by increasing amounts of Lck, whereas increases of CD45 density had the opposing effect. Furthermore, binding of the ZAP70 SH2 domains was shown to protect phosphorylated ζ from CD45-mediated dephosphorylation, whereas Lck-ζ co-clustering led to intense ITAM phosphorylation even at the presence of very high CD45 concentrations. These data are consistent with our observations of CD45, pY142-ζ and pZAPY493 co-existence within the Rab11-decorated structures under conditions of Lck overexpression and can be projected to a model in which CD45 maintains basal phosphorylation of endosomal ζ. Upon T cell activation, the documented CD45-Lck time-dependent co-clustering and segregation at sites of TCR engagement (37) could counteract the dominance/kinetic advantage of CD45. In parallel, the time-dependent enrichment of Lck and ZAP70 to the EC (as has been shown in CD2-triggered T cells (38)) could serve the same purpose, with the end result being the creation of a depository of activated signaling mediators supplied to the IS.

Concerning the parameters governing Lck and ζ chain EC residency, we showed that the colocalization scores between ζ and Rab11 were invariably high in unstimulated human T cells and T cell lines but also in B cell lines where ζ chain was ectopically expressed. Notably, the presence of endosomal ζ was independent of SFK activity or TCR expression at the cell surface, highlighting that although ITAM phosphorylation or the detachment of “loosely-attached” ζ from the TCR complex are important within the broader and heterogeneous landscape of ζ chain internalization (13–15, 20), they do not constitute primary mechanisms promoting its residency within Rab11 endosomes.

In parallel, we identified two decisive parameters for Lck internalization. The first one relies within intrinsic characteristics of its SH4 domain. The chimeric Fyn-Lck and Lyn-Lck differ from the WT protein only in a short (11 and13 amino acids, respectively) and unstructured N-terminal stretch. Although all three different SH4s undergo myristoylation and palmitoylation and are mediating SFK anchoring at the inner leaflet of the PM, the divergency in their sequences (26) sufficed to dramatically alter the subcellular distribution of Lck, with the two chimeric forms failing to reside within the Rab11^+^EC. The recognition of SFK SH4 domains as a crucial structural element for endosomal residency and the correlation with Lck signaling efficacy reveals a previously unrecognized aspect of SH4’s importance in SFK function. This finding may also explain in part Lck ‘s optimal adaptation for the TCR signaling apparatus compared to other antigen receptors (31).

The second key parameter for Lck (and CD45) colocalization with Rab11 is the presence of the TCRζ chain. Ectopically expressed Lck in BJAB-naturally devoid of ζ-exhibited exclusive PM residency, a pattern also shared by CD45. Introduction of TCRζ in this B cell environment led to the redistribution of both Lck and CD45 to the Rab11^+^ EC. Accordingly, endosomal Lck and CD45 were diminished in jurkat cells where the ζ chain was silenced whereas, endosomal CD45 was undetectable in non-T cell PBMCs. These observations indicate that Lck and CD45 transportation to the Rab11 compartment is driven by and occurs in parallel to the ζ chain. This prompts the question of what underlying mechanism prohibits the Lck SH4 chimeras from colocalizing with Rab11 in a cellular context where the ζ chain is sufficiently expressed. In a previous study (24), we showed that the lateral organization of membrane-resident proteins is governed by their residency within compatible (‘like’) or incompatible (‘unlike’) lipid boundary environments, a property that respectively decides their spatial proximity or segregation. Based on this, a plausible explanation for the failure of Fyn-Lck and Lyn-Lck to internalize, is that the biophysical characteristics of their SH4 domains hinder their spatial juxtaposition to the ζ chain and thus their co-transportation to the Rab11^+^EC. However, the SH4 chimeric forms do gain access to ζ molecules that are incorporated within the TCR complex, as evidenced by pY142-ζ induction upon anti-CD3 stimulation (which specifically engages surface TCRs). We were therefore prompted to investigate whether TCR triggering could alter the subcellular distribution of PM-resident Fyn-Lck and Lyn-Lck. TCR internalization alongside ζ dissociation from the complex has been demonstrated for receptors engaged both by pMHC and also after incubation with a-CD3 antibodies (14–16) and Fig.S3F). If Lck is co-transported to the Rab11^+^ compartment with “loosely attached” ζ chain that disengages from the TCR, then Fyn-Lck and Lyn-Lck should accumulate at the EC in stimulated cells. On the contrary though, Fyn-Lck and Lyn-Lck remained at the PM following TCR engagement and could not be detected at the Rab11^+^ compartment, suggesting that Lck is not co-internalizing with the fraction of ζ molecules detaching from the receptor complex. Furthermore, we could not detect enrichment of pY142-ζ in a-CD3 stimulated Fyn-Lck and Lyn-Lck cell lines, despite the presence of total ζ at this location, signifying that the Rab11^+^ ζ pool is not derived from the immediate internalization of the fraction that disengages from the TCR complex. This is in agreement with the work of Del Río-Iñiguez et al. (39) demonstrating that CD3/CD28 crosslinking failed to induce endosomal enrichment of Lck and pZAP during the time interval that ζ and ZAP phosphorylation, TCR/CD3 internalization and ζ dissociation from the receptor complex are known to occur. Rather, they showed that TCR engagement diminished the intensity of the Lck-containing endosomal compartment over time, suggesting translocation of the Rab11 co-resident molecules to the plasma membrane rather than their endosomal accumulation during early activation.

In sharp contrast to a-CD3 stimulation, accumulation of Fyn-Lck and Lyn-Lck in Rab11^+^ endosomes was observed in pervanadate-treated cells, which coincided with pY142-ζ within the compartment. Nevertheless, pervanadate treatment per se did not suffice to induce Lck internalization in the absence of the ζ chain. The pervanadate-induced appearance of endosomal Fyn-Lck and Lyn-Lck implies that under this extremely robust, non-physiological type of activation SH4-imposed restrictions can be alleviated so that the chimeric proteins acquire vicinity to, the otherwise inaccessible, ζ fraction driving Lck internalization. This is a perfectly plausible scenario, since pervanadate treatment has been shown to induce changes in diffusion rates and protein aggregation profiles (40,41).

Collectively our data show that endosomal Lck originates from the PM, at sites where its juxtaposition to the ζ chain enables their co-trafficking to the Rab11^+^compartment. However, the ζ species driving Lck internalization are distinct from the “loosely attached” fraction previously described as recycling to and from the TCR complex (13), supporting the existence of two distinct ζ chain populations at the PM: one incorporated within the TCR, and another residing at the plasma membrane as autonomous units, which we hereafter refer to as “standalone ζ”.

The capacity of the ζ chain to acquire self-directed plasma membrane residency has been documented in cell lines and hybridomas lacking TCR surface expression (13) and Figs.2A, 2E). Furthermore, La Gruta et al. (14) have reported that in unstimulated T cell hybridomas with fully assembled TCRs, a small proportion of PM-localized ζ existed uncoupled from the rest of the receptor complex and that this fraction was significantly upregulated following antigen stimulation. Alongside our findings two scenarios can be envisioned for the origin of standalone ζ: a) a fraction of the newly synthesized protein may be delivered at the PM as an independent unit uncoupled from the TCR. However, this possibility is challenged by evidence that ζ is the limiting factor for TCR assembly and surface expression, with a synthesis rate estimated to be about one-tenth that of the other chains (33). Moreover, induced increases in ζ synthesis were directly proportional to increases in surface TCR expression, indicating that its vast majority was incorporated in the receptor complex (32) and b) a subset of the “loosely attached” ζ molecules that constitutively recycle to and from the surface TCRs (13), instead of directly internalizing, may be laterally mobilized at the plasma membrane to form the standalone pool. The TCR ligation-mediated enrichment of the standalone ζ pool (14) could thus be mediated via delivery of the Rab11^+^endosomal molecular assembly towards the PM (6,17,19) and/or by augmented disengagement of ζ from ligated receptor complexes (14,15).

It is widely accepted that TCR signaling is orchestrated by spatiotemporally regulated vesicular trafficking. TCR ligation has been shown to promote polarization of vesicles containing Lck, phosphorylated ζ, ZAP-70, LAT etc towards the immunological synapse, a mechanism suggested to provide rapid and sustained supply of signalling mediators at sites of receptor engagement (3,5,6). Within this context, we propose the following model shaped by our findings: At the PM, ζ can be found in two distinct pools, incorporated within the TCR complex and as a standalone population. The latter constitutes the driving force for Lck co-internalization into the Rab11^+^ EC and formation of signalling assemblies that are maintained at steady-state phosphorylation by the dominance of CD45. Upon receptor ligation these assemblies are mobilized towards signalling “hot-spots” where increased ζ phosphorylation and subsequent ZAP-70 recruitment are driven by local Lck enrichment and/or CD45 segregation.

Upon vesicular release, phosphorylated ζ and its co-residents could either integrate at the PM as autonomous units or get incorporated in TCR complexes (both unligated and engaged) to replenish the loss of disengaged ζ. In either case, they establish a foundation of pre-primed signalling platforms that enable rapid and sustained propagation of downstream signals, decreasing the dependence on continuous receptor engagement. This mechanism complements the conceptual framework regarding the remarkable T cell sensitivity to low-affinity and sparse density of pMHC complexes, whereas it can provide an advancement on our understanding of responses associated with bystander, non-engaged TCRs (10–12,42)

Furthermore, our findings on a previously unidentified mechanism of Lck and ζ co-trafficking, become particularly relevant within the context of cell therapies with Chimeric Antigen Receptor (CARs). CAR-T cells heavily rely on fine-tuned Lck function for optimal effectiveness and safety profiles (43). Both hypeactivation or underperformance of CAR signalling can compromise clinical outcomes either by inducing adverse toxicity or by weakening therapeutic efficacy respectively. Understanding and substantiating the operation of distinct ζ pools and their contribution in shaping T cell activation thresholds and signalling potency, independently of sustained TCR or CAR engagement, could become the base of strategies for enhance CAR-T sensitivity, durability, performance and safety.

## Author Contributions

Conceptualization: KN

Methodology: KK, NK, AK, VR, KN,

Investigation: KK, NK, IT, ET, AK

Visualization: KN, KK, NK,

Supervision: KN

Writing: original draft: KN

Writing: review & editing: KN, KK, AK, VR

## Acknowledgments

We are grateful to Prof. Björn Lillemeier (Faculty of Biology, University of Freiburg) for provision of the J76 cell line and to Oreste Acuto (University of Oxford) for provision of Lck constructs and valuable feedback on the manuscript.

## Materials and methods

### Antibodies and Reagents

Rabbit anti-pSrc (Y416), detecting Lck pY394 (Cat: 2101), rabbit anti-Lck (73A5) (Cat: 2787), rabbit phospho-Syk (Tyr525/526), detecting Zap-70 pY493 (Cat:2711), rabbit phospho-CD79a (Tyr182) (Cat:5173) and rabbit anti-Rab11 (D4F5) (Cat: 5589) were purchased from Cell Signaling Technology (CST). Mouse anti-Lck-PE (3A5) (Cat: sc-433), mouse anti-Lck (3A5) (unconjugated) (Cat: sc-433) and rat anti-CD45 (YAML 501.4) (Cat: sc-65344) were purchased from Santa Cruz Biotechnology. Rabbit polyclonal anti-Lck (Cat: 85804), used for confocal microscopy, was purchased from Novus Biologicals. Rabbit anti-CD3 zeta (phospho Y142) [EP265(2)Y] (ab68235), used for confocal microscopy, was from abcam. Mouse anti-CD247 (pY142) Alexa Fluor 647 (Cat: 558489) and mouse anti-CD247 (pY142) (Cat: 558402) were from BD Biosciences. Chicken anti-Green Fluorescent Protein (GFP) (GFP-1010) was from Aves Labs. Mouse anti-human CD3 antibody (clone UCHT1) Alexa Fluor 647 (Cat: 300416), used for surface staining in FACS and mouse purified anti-human CD3 (clone UCHT1, Cat: 300402), used for stimulation assays, as well as for confocal microscopy, were purchased from BioLegend. Secondary antibodies for confocal microscopy and FACS goat anti-mouse IgG Alexa Fluor 568 (Cat: A11001), goat anti-rabbit IgG Alexa Fluor 647 (Cat: A21245) and goat anti-rat IgG Alexa Fluor 568 (Cat: A11077) were from Thermo Fischer Scientific. The pan-SFK inhibitor PP2 was from Sigma Aldrich (529573).

### Cells, treatments and stimulations

All cell lines were cultured at 37 C in a humidified incubator (PHCbi) with 5% CO_2_. Human embryonic kidney (HEK) 293T cells (ATCC® CRL-3216) were cultured in DMEM (Gibco) supplemented with 10% fetal bovine serum (Gibco). Jurkat Clone 20 (C20), JCaM1.6, J76 (jurkat subline devoid of endogenous TCR alpha and beta chains) and BJAB, were maintained in RPMI 1640 (Gibco) containing 10% FBS. Cells engineered for Dox-inducible expression were cultured in RPMI 1640 supplemented with 10% tetracycline-free FBS. The BJAB-LckWT cell line, expressing wild type human Lck tagged with GFP in its C-terminus has been previously described (31). All lines were routinely tested for mycoplasma (Universal Mycoplasma Detection Kit-ATCC, 30-1-12K), lineage-specific surface markers and CD45 expression.

Peripheral blood mononuclear cells (PBMCs) were separated from whole blood of healthy donors by density gradient centrifugation (Ficoll-Paque™ PLUS, Cytiva/GE Healthcare). Primary human CD4^⁺^ T cells were purified from PBMCs using the Untouched™ Human CD4 kit (Thermo Fisher Scientific, Cat: 11346D), purity: ≥95%. Cells were routinely kept in culture (RPMI 1640, 10% FBS) overnight prior to usage.

For signal transduction triggering experiments, cells (1×10^6^ cells/mL) were stimulated by 1 µg/mL soluble anti–human CD3ε (clone UCHT1) for 2min at 37 °C. For subcellular distribution and colocalization evaluations by confocal imaging, cells were first activated in an Eppendorf tube with 1 µg/mL anti–CD3ε for 1min at 37 °C and then transferred onto poly-D-lysine–coated coverslips and allowed to adhere for an additional 10 minutes at 37 °C before fixation and staining.

For pan-SFK inhibition, cells were cultured with 30μM PP2 for 18h in the humidified incubator. Pervanadate solutions were prepared immediately prior to each assay to ensure maximal activity. For pervanadate stimulation, cells, plated on coverslips, were incubated with 100μM pervanadate for 2, 5 and 10min at 37 °C (depending on the experiment, as indicated in respective figure legends) and immediately fixed to preserve phospho-epitopes and endosomal architecture.

### Cloning and plasmids

The cDNAs of human LckWT, the LckY394F, ΔSH4 mutants, and human TCRζ chain (cloned in the lentiviral vector pHR-Bin) were kindly provided by O. Acuto (University of Oxford). Fyn-Lck and Lyn-Lck were generated by PCR using LckΔSH4 as template, and oligonucleotides encoding the SH4 domains of Fyn (amino acids 1–11, Forward primer: ATAAGAATGCGGCCGCATGGGCTGTGTGCAATGTAAGGATAAAGAAGCAGATGACTGGATGGAAAA CATC) and Lyn (amino acids 1–13, Forward primer: ATAAGAATGCGGCCGCATGGGATGTATAAAATCAAAAGGGAAAGACAGCTTGAGTGATGACTGGATGGAAAACATC) respectively. All Lck constructs were cloned in the pLVX-Tight-Puro vector (Clontech Laboratories), between 5’ NotI and 3’ EcoRI restriction sites. The lentiviral packaging plasmids pVSVG (Plasmid #8454) and pSPAX2 (Plasmid #12260) were purchased from Addgene.

### Generation of Dox-inducible cell lines

Lentiviral particles were generated using HEK 293T cells as the packaging system. Cells were seeded one day prior to transfection and used at 60–80% confluency. Transfection was performed using polyethylenimine (PEI, Polysciences) following a standard PEI-based protocol. The transfer plasmids pLVX-Tet-On-Advanced and pLVX-Tight-Puro (Clontech Laboratories), encoding the rtTA-Advanced transactivator and genes of interest, respectively, were co-transfected with the packaging plasmids pSPAX2 and pVSVG. DNA and PEI were diluted in 200 µL of Opti-MEM (Gibco, incubated at room temperature for 20 minutes, and added dropwise to the cells. Culture medium was replaced with fresh DMEM containing 10% FBS after 18 hours. Viral supernatants were collected 48 hours after transfection and used for transduction.

For transduction, approximately, 5×10^5^ cells were incubated with viral supernatants for 24 hours in the presence of 5µg/mL hexadimethrine bromide (Polybrene; Sigma-Aldrich). After incubation, the viral media were discarded and replaced with fresh RPMI supplemented with 10% FBS. Cells were allowed to recover for 48 hours at 37 °C and 5% CO_2_, after which selection was initiated using 5 µg/mL puromycin and 200 µg/mL Geneticin (G418). For induced gene of interest expression, cells were cultured in the presence of 1µg/mL doxycycline (Sigma-Aldrich) for 24h prior to analyses.

### CRISPR/Ca9-mediated knockout of TCRζ

Biallelic TCRζ-inactivated clones, were generated using the lentiviral vector lentiCRISPR v2 (Addgene plasmid # 52961), which co-expresses a human codon optimized Cas9 with a nuclear localization signal, under the EF-1a immediate early promoter, together with a gRNA driven by the U6 promoter. A previously validated gRNA targeting exon 2 of the *CD247* gene *(44)* (sequence: AGCAGAGTTTGGGATCCAGC) was inserted downstream of the U6 promoter sequence. HEK293 cells were co-transfected with the lentiCRISPR v2-gRNA-TCRζ lentiviral vector and pMD2.G (Addgene plasmid #12259), psPAX2 (Addgene Plasmid #12260). Jurkat cells were transduced with viral supernatants as described above, after 24h, 1ml of fresh medium was added, and the following day the cells were subjected to puromycin selection. ζ knockout was confirmed by FACS analysis and with a-CD3ε to confirm downregulation of surface TCR expression

### Flow cytometry

For intracellular staining, cells were fixed in BD Phosflow™ Fix Buffer I (BD Biosciences) at 37 °C for 10 minutes. Fixed cells were centrifuged and permeabilized in 150 µl of PBS supplemented with 0.5% BSA and 0.5% Triton X-100 (Millipore) for 15 minutes at 37 °C. Cells were then incubated with primary antibodies, diluted in PBS with 0.5% BSA and 0.1% Tween 20, for 1 hour at 37 °C. Following two washes, where needed, samples were incubated with the corresponding fluorescent secondary antibodies for 45 minutes at 37 °C. Cells were washed twice and then analyzed on BD Accuri™ C6 Plus Flow Cytometer.

For TCR downregulation estimations after pervanadate treatment, cells were resuspended in 100µl of FACS buffer (PBS containing 0.5% BSA) and incubated on ice for 15 minutes to block nonspecific binding. After centrifugation, supernatants were removed, and cells were stained with alexa 647-conjugated CD3ε diluted in FACS buffer for 30 minutes on ice. Acquired data were analyzed by FlowJo Software (BD Biosciences).

### Immunofluorescence, confocal microscopy image acquisition and analysis

For the preparation of confocal microscopy samples, single‐cell suspensions were seeded onto poly-D-lysine (Gibco)–coated coverslips (Marienfeld Superior). Cells were fixed with BD Phosflow™ Fix Buffer I (BD Biosciences) for 10 minutes at 37 °C, rinsed once in DPBS (with Ca^2⁺^/Mg^2⁺^), and permeabilized for 5min at 37 °C in DPBS containing 1% BSA and 0.1% Triton X-100. After a DPBS wash, nonspecific sites were blocked by incubation in DPBS supplemented with 1% BSA and 0.1% Tween 20 (blocking buffer) for 10 minutes at 37 °C. Samples were incubated with the indicated primary antibodies, diluted in Blocking/Wash Buffer for 1h at 37°C, washed three times and then incubated for 1h at 37°C with the corresponding fluorescent-conjugated secondary antibodies, diluted in blocking/wash buffer. After three final washes, coverslips were mounted on microscope slides using ProLong® Gold Antifade Mountant (Thermo Fisher Scientific).

Images were captured on a Leica TCS SP5 confocal microscope (Leica Microsystems) using a 63×/1.4 NA oil-immersion objective. Fluorophores were excited at 488 nm, 561 nm, and 633 nm, with z-stacks collected at 0.8 μm intervals. All raw datasets were processed in ImageJ: Plasma membrane/Endoplasmic compartment ratios: ROIs delineating the PM and Endosomal compartment areas were defined by CD45 and Rab11 staining respectively, for each cell, and mean fluorescence intensities were measured within the corresponding compartments to compute the ratio. Colocalization analyses: Pearson’s correlation coefficients were determined using the Coloc2 plugin of Imaje J. For each cell, a single midplane optical section was selected, and ROIs were defined as indicated in corresponding figure legends

## Statistical analysis

Statistical analysis was performed using GraphPad Prism 9 (GraphPad Software). Data were analyzed using unpaired two-tailed Student t test or multiple unpaired two-tailed Student t test. All data are presented as mean +/- SD. A p-value less than 0.05 was considered significant (*P < 0.05, **P < 0.01, ***P < 0.001, ****P < 0.0001; ns: not significant). All experiments were repeated with sufficient reproducibility.

## References

1. Acuto O, Bartolo V Di, Michel F. Tailoring T-cell receptor signals by proximal negative feedback mechanisms. Nat Rev Immunol (2008) 8:699–712. doi: 10.1038/nri2397

2. Monks CRF, Freiberg BA, Kupfer H, Sciaky N, Kupfer A. Three-dimensional segregation of supramolecular activation clusters in T cells. Nature (1998) 395:82–86. doi: 10.1038/25764

3. Balagopalan L, Barr VA, Samelson LE. Endocytic events in TCR signaling: focus on adapters in microclusters. Immunol Rev (2009) 232:84–98. doi: 10.1111/j.1600-065X.2009.00840.x

4. Evnouchidou I, Caillens V, Koumantou D, Saveanu L. The role of endocytic trafficking in antigen T cell receptor activation. Biomed J (2022) 45:310–320. doi: 10.1016/j.bj.2021.09.004

5. Das V, Nal B, Dujeancourt A, Thoulouze M-I, Galli T, Roux P, Dautry-Varsat A, Alcover A. Activation-induced polarized recycling targets T cell antigen receptors to the immunological synapse; involvement of SNARE complexes. Immunity (2004) 20:577–88. doi: 10.1016/s1074-7613(04)00106-2

6. Soares H, Henriques R, Sachse M, Ventimiglia L, Alonso MA, Zimmer C, Thoulouze M-I, Alcover A. Regulated vesicle fusion generates signaling nanoterritories that control T cell activation at the immunological synapse. Journal of Experimental Medicine (2013) 210:2415–2433. doi: 10.1084/jem.20130150

7. Liu H, Rhodes M, Wiest DL, Vignali DAA. On the Dynamics of TCR:CD3 Complex Cell Surface Expression and Downmodulation. Immunity (2000) 13:665–675. doi: 10.1016/S1074-7613(00)00066-2

8. Valitutti S, Müller S, Salio M, Lanzavecchia A. Degradation of T Cell Receptor (TCR)–CD3-ζ Complexes after Antigenic Stimulation. J Exp Med (1997) 185:1859–1864. doi: 10.1084/jem.185.10.1859

9. Niedergang F, Dautry-Varsat A, Alcover A. Peptide antigen or superantigen-induced down-regulation of TCRs involves both stimulated and unstimulated receptors. J Immunol (1997) 159:1703–10.

10. José ES, Borroto A, Niedergang F, Alcover A, Alarcón B. Triggering the TCR Complex Causes the Downregulation of Nonengaged Receptors by a Signal Transduction-Dependent Mechanism. Immunity (2000) 12:161–170. doi: 10.1016/S1074-7613(00)80169-7

11. Monjas A, Alcover A, Alarcón B. Engaged and Bystander T Cell Receptors Are Down-modulated by Different Endocytotic Pathways. Journal of Biological Chemistry (2004) 279:55376–55384. doi: 10.1074/jbc.M409342200

12. Alcover A, Alarcón B, Di Bartolo V. Cell Biology of T Cell Receptor Expression and Regulation. Annu Rev Immunol (2018) 36:103–125. doi: 10.1146/annurev-immunol-042617-053429

13. Ono S, Ohno H, Salto T. Rapid turnover of the CD3ζ chain independent of the TCR-CD3 complex in normal T cells. Immunity (1995) 2:639–644. doi: 10.1016/1074-7613(95)90008-X

14. La Gruta NL, Liu H, Dilioglou S, Rhodes M, Wiest DL, Vignali DAA. Architectural Changes in the TCR:CD3 Complex Induced by MHC:Peptide Ligation. The Journal of Immunology (2004) 172:3662–3669. doi: 10.4049/jimmunol.172.6.3662

15. Lanz A-L, Masi G, Porciello N, Cohnen A, Cipria D, Prakaash D, Bálint Š, Raggiaschi R, Galgano D, Cole DK, et al. Allosteric activation of T cell antigen receptor signaling by quaternary structure relaxation. Cell Rep (2021) 36:109375. doi: 10.1016/j.celrep.2021.109375

16. Brazin KN, Mallis RJ, Boeszoermenyi A, Feng Y, Yoshizawa A, Reche PA, Kaur P, Bi K, Hussey RE, Duke-Cohan JS, et al. The T Cell Antigen Receptor α Transmembrane Domain Coordinates Triggering through Regulation of Bilayer Immersion and CD3 Subunit Associations. Immunity (2018) 49:829-841.e6. doi: 10.1016/j.immuni.2018.09.007

17. Yudushkin IA, Vale RD. Imaging T-cell receptor activation reveals accumulation of tyrosine-phosphorylated CD3ζ in the endosomal compartment. Proceedings of the National Academy of Sciences (2010) 107:22128–22133. doi: 10.1073/pnas.1016388108

18. Redpath GMI, Ecker M, Kapoor-Kaushik N, Vartoukian H, Carnell M, Kempe D, Biro M, Ariotti N, Rossy J. Flotillins promote T cell receptor sorting through a fast Rab5–Rab11 endocytic recycling axis. Nat Commun (2019) 10:4392. doi: 10.1038/s41467-019-12352-w

19. Evnouchidou I, Chappert P, Benadda S, Zucchetti A, Weimershaus M, Bens M, Caillens V, Koumantou D, Lotersztajn S, van Endert P, et al. IRAP-dependent endosomal T cell receptor signalling is essential for T cell responses. Nat Commun (2020) 11:2779. doi: 10.1038/s41467-020-16471-7

20. Bouchet J, del Río-Iñiguez I, Vázquez-Chávez E, Lasserre R, Agüera-González S, Cuche C, McCaffrey MW, Di Bartolo V, Alcover A. Rab11-FIP3 Regulation of Lck Endosomal Traffic Controls TCR Signal Transduction. The Journal of Immunology (2017) 198:2967–2978. doi: 10.4049/jimmunol.1600671

21. Gorska MM, Stafford SJ, Cen O, Sur S, Alam R. Unc119, a Novel Activator of Lck/Fyn, Is Essential for T Cell Activation. J Exp Med (2004) 199:369–379. doi: 10.1084/jem.20030589

22. Kabouridis PS. S-acylation of LCK protein tyrosine kinase is essential for its signalling function in T lymphocytes. EMBO J (1997) 16:4983–4998. doi: 10.1093/emboj/16.16.4983

23. Resh MD. Myristylation and palmitylation of Src family members: The fats of the matter. Cell (1994) 76:411–413. doi: 10.1016/0092-8674(94)90104-X

24. Porciello N, Cipria D, Masi G, Lanz A-L, Milanetti E, Grottesi A, Howie D, Cobbold SP, Schermelleh L, He H-T, et al. Role of the membrane anchor in the regulation of Lck activity. Journal of Biological Chemistry (2022) 298:102663. doi: 10.1016/j.jbc.2022.102663

25. Owen DM, Rentero C, Rossy J, Magenau A, Williamson D, Rodriguez M, Gaus K. PALM imaging and cluster analysis of protein heterogeneity at the cell surface. J Biophotonics (2010) 3:446–454. doi: 10.1002/jbio.200900089

26. Eshaq AM, Flanagan TW, Hassan S-Y, Al Asheikh SA, Al-Amoudi WA, Santourlidis S, Hassan S-L, Alamodi MO, Bendhack ML, Alamodi MO, et al. Non-Receptor Tyrosine Kinases: Their Structure and Mechanistic Role in Tumor Progression and Resistance. Cancers (Basel) (2024) 16:2754. doi: 10.3390/cancers16152754

27. Abraham KM, Levin SD, Marth JD, Forbush KA, Perlmutter RM. Thymic tumorigenesis induced by overexpression of p56lck. Proceedings of the National Academy of Sciences (1991) 88:3977–3981. doi: 10.1073/pnas.88.9.3977

28. Luo K, Sefton BM. Activated lek Tyrosine Protein Kinase Stimulates AntigenIndependent Interleukin-2 Production in T Cells. Mol Cell Biol (1992) 12:4724–4732. doi: 10.1128/mcb.12.10.4724-4732.1992

29. Manz BN, Tan YX, Courtney AH, Rutaganira F, Palmer E, Shokat KM, Weiss A. Small molecule inhibition of Csk alters affinity recognition by T cells. Elife (2015) 4: doi: 10.7554/eLife.08088

30. Crotzer VL, Mabardy AS, Weiss A, Brodsky FM. T Cell Receptor Engagement Leads to Phosphorylation of Clathrin Heavy Chain during Receptor Internalization. J Exp Med (2004) 199:981–991. doi: 10.1084/jem.20031105

31. Koutras N, Morfos V, Konnaris K, Kouvela A, Shaukat A-N, Stathopoulos C, Stamatopoulou V, Nika K. Integrated signaling and transcriptome analysis reveals Src family kinase individualities and novel pathways controlled by their constitutive activity. Front Immunol (2023) 14: doi: 10.3389/fimmu.2023.1224520

32. Weissman AM, Frank SJ, Orloff DG, Merćep M, Ashwell JD, Klausner RD. Role of the zeta chain in the expression of the T cell antigen receptor: genetic reconstitution studies. EMBO J (1989) 8:3651–3656. doi: 10.1002/j.1460-2075.1989.tb08539.x

33. Minami Y, Weissman AM, Samelson LE, Klausner RD. Building a multichain receptor: synthesis, degradation, and assembly of the T-cell antigen receptor. Proceedings of the National Academy of Sciences (1987) 84:2688–2692. doi: 10.1073/pnas.84.9.2688

34. Dustin ML. Recent advances in understanding TCR signaling: a synaptic perspective. Fac Rev (2023) 12: doi: 10.12703/r/12-25

35. Marie‐Cardine A, Maridonneau‐Parini I, Fischer S. Activation and internalization of p56 ^lck^upon CD45 triggering of Jurkat cells. Eur J Immunol (1994) 24:1255–1261. doi: 10.1002/eji.1830240603

36. Hui E, Vale RD. In vitro membrane reconstitution of the T-cell receptor proximal signaling network. Nat Struct Mol Biol (2014) 21:133–142. doi: 10.1038/nsmb.2762

37. Freiberg BA, Kupfer H, Maslanik W, Delli J, Kappler J, Zaller DM, Kupfer A. Staging and resetting T cell activation in SMACs. Nat Immunol (2002) 3:911–917. doi: 10.1038/ni836

38. Marie-Cardine A, Fischer S, Gorvel J-P, Maridonneau-Parini I. Recruitment of Activated p56 on Endosomes of CD2-triggered T Cells, Colocalization with ZAP-70. Journal of Biological Chemistry (1996) 271:20734–20739. doi: 10.1074/jbc.271.34.20734

39. del Río-Iñiguez I, Vázquez-Chávez E, Cuche C, Di Bartolo V, Bouchet J, Alcover A. HIV-1 Nef Hijacks Lck and Rac1 Endosomal Traffic To Dually Modulate Signaling-Mediated and Actin Cytoskeleton–Mediated T Cell Functions. The Journal of Immunology (2018) 201:2624–2640. doi: 10.4049/jimmunol.1800372

40. Bennett PA, Dixon RJ, Kellie S. The phosphotyrosine phosphatase inhibitor vanadyl hydroperoxide induces morphological alterations, cytoskeletal rearrangements and increased adhesiveness in rat neutrophil leucocytes. J Cell Sci (1993) 106:891–901. doi: 10.1242/jcs.106.3.891

41. Park J, Hill MM, Hess D, Brazil DP, Hofsteenge J, Hemmings BA. Identification of Tyrosine Phosphorylation Sites on 3-Phosphoinositide-dependent Protein Kinase-1 and Their Role in Regulating Kinase Activity. Journal of Biological Chemistry (2001) 276:37459–37471. doi: 10.1074/jbc.M105916200

42. Nieves DJ, Pandzic E, Gunasinghe SD, Goyette J, Owen DM, Justin Gooding J, Gaus K. The T cell receptor displays lateral signal propagation involving non-engaged receptors. Nanoscale (2022) 14:3513–3526. doi: 10.1039/D1NR05855J

43. Sun C, Shou P, Du H, Hirabayashi K, Chen Y, Herring LE, Ahn S, Xu Y, Suzuki K, Li G, et al. THEMIS-SHP1 Recruitment by 4-1BB Tunes LCK-Mediated Priming of Chimeric Antigen Receptor-Redirected T Cells. Cancer Cell (2020) 37:216-225.e6. doi: 10.1016/j.ccell.2019.12.014

44. Hintz HM, Snyder KM, Wu J, Hullsiek R, Dahlvang JD, Hart GT, Walcheck B, LeBeau AM. Simultaneous Engagement of Tumor and Stroma Targeting Antibodies by Engineered NK-92 Cells Expressing CD64 Controls Prostate Cancer Growth. Cancer Immunol Res (2021) 9:1270–1282. doi: 10.1158/2326-6066.CIR-21-0178

