## Supplemental Information for "TCRζ-Driven Pre-Signaling Organization of Lck in Rab11^+^ Endosomes Shapes TCR Activation"

### Supplemental Figures

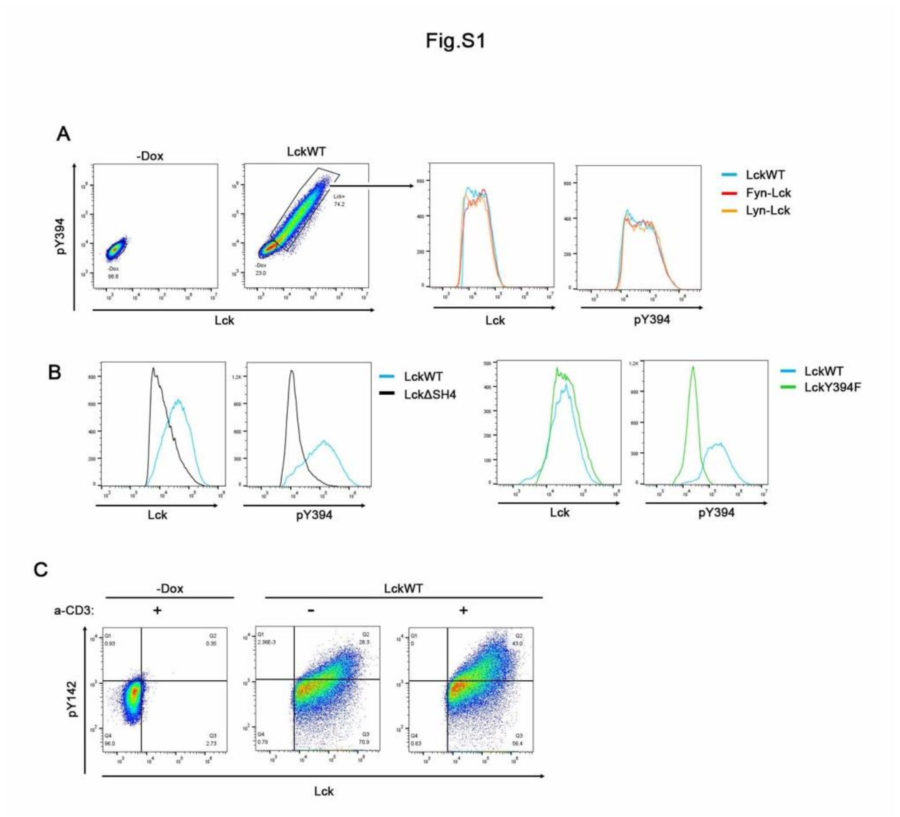


**Figure S1. (Related to Fig.1).**

A. Gating strategy for FACS analyses and characterization of the cell lines used in this study. JcaM1.6-LckWT cells and their -Dox counterparts were stained for Lck and pY394 and analysed by FACS. Single cells were selected from FSC-A versus FGC-H dot plots (not shown). Lck-expressing populations were isolated based on gating set by respective antibodies fluorescence in the -Dox samples. MFIs for Lck and pY394 were calculated within the Lck$^{+}$ subsets. Representative overlay histograms for Lck and pY394 of LckWT and the SH4 chimeras are shown on the right. This gating strategy was followed for all subsequent FACS-based analyses of Dox-inducible cell lines in this study.

B. The LckΔSH4 and LckY394F mutants have compromised Lck activity. Representative overlay histograms for total protein (Lck) expression and pY394 levels of LckΔSH4 and LckY394F, compared to the WT protein.

C. Gating strategy for a-CD3 stimulation assays. LckWT cells and their -Dox counterparts were either left untreated or stimulated with anti-CD3 for 2min at 37$^{o}$C and stained for Lck and pY142-ζ. Initial gating steps as in (A), Lck $^{+}$ subsets were further gated according to anti-pY142 fluorescence level in the -Dox samples. Graphs in Fig.1C , display percentages of cells in the upper right quadrant, representative of the pY142⁺ populations (or pY493⁺ of ZAP 70, as indicated).

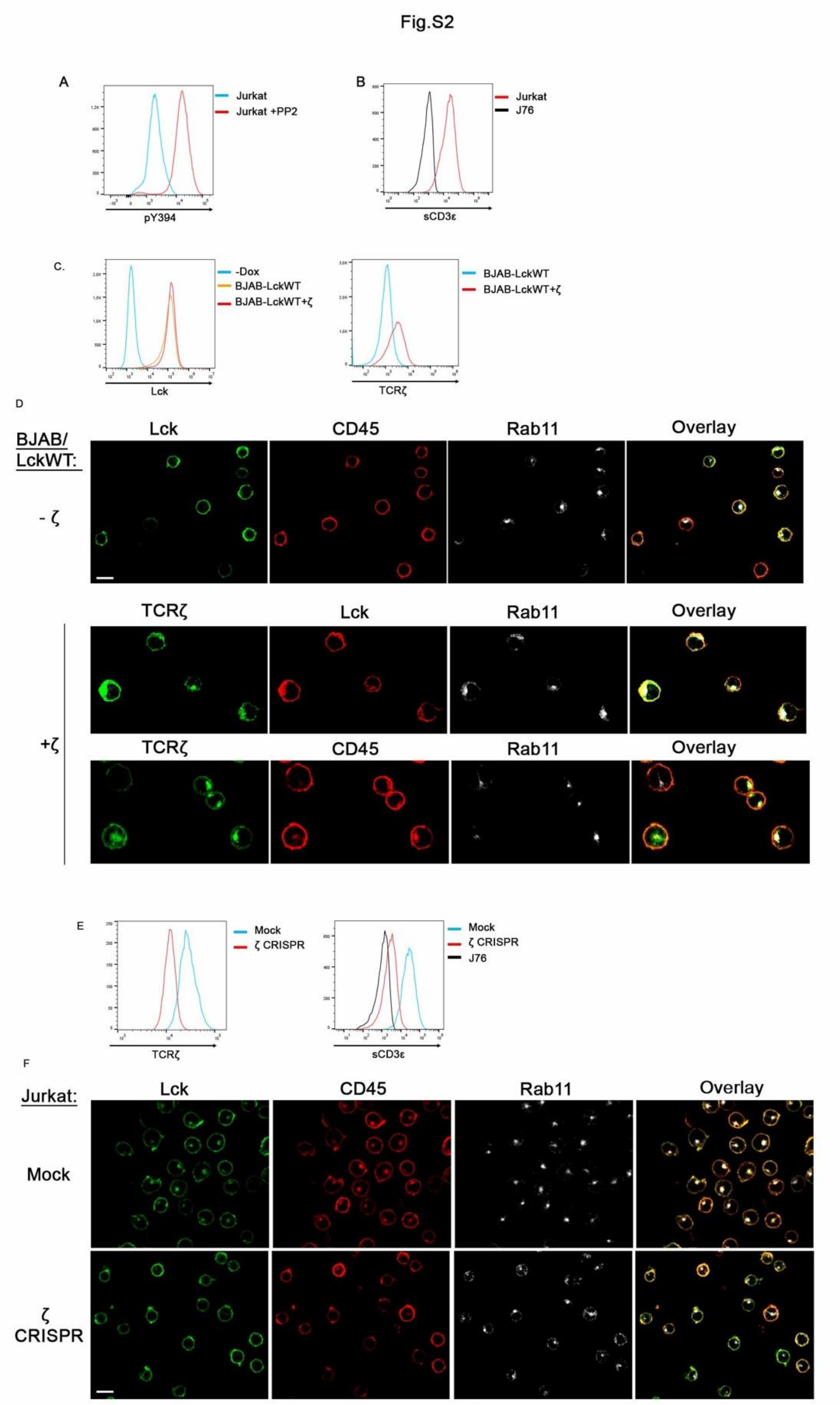


**<u>Figure S2 (Related to Fig,2)</u>**

A. Verification of PP2-mediated diminished SFK activity in Jurkat cells. Representative overlay histogram of Jurkat cells cultured for 18h in the absence or presence of 30μM PP2, stained with a-pY394 and analysed by FACS

B. Phenotypic verification of the J76 lines. Overlay histograms of J76 and Jurkat cells (serving as a positive marker for surface TCR expression) stained for surface CD3 (sCD3).

C. Characterization of the BJAB-LckWT lines transduced with TCRζ. *Left*: Overlay histogram of BJAB-LckWT cells cultured in the presence or absence of Dox (-Dox) and stained with a-Lck. *Right*: BJAB-LckWT cells and their counterparts transduced with TCRζ stained a-TCRζ. Scale bar, 10μm

D. Low-magnification images of BJAB samples, corresponding to Fig. 2E, to allow visualization of localization patterns across a broader cell population. Due to antibody species restrictions images taken from BJAB-LckWT cells (31) co-expressing TCRζ (Fig.2E, and here), Lck was visualized in the corresponding panels by anti-GFP antibody raised in chicken.

E. Phenotypic verification of the ζ-CRISPR line. Representative overlay histograms for the ζ-CRISPR jurkat cells and their mock-transduced counterparts stained for TCRζ (*left histogram*) and surface CD3 (sCD3ε) (*right histogram*) expression, respective sCD3 staining in J76 serves as a positive control for TCR downregulation.

F. Low-magnification images of Mock-transduced or ζ-CRISPR cells, corresponding to Fig.2F. Scale bars, 10μm.

Fig.S3

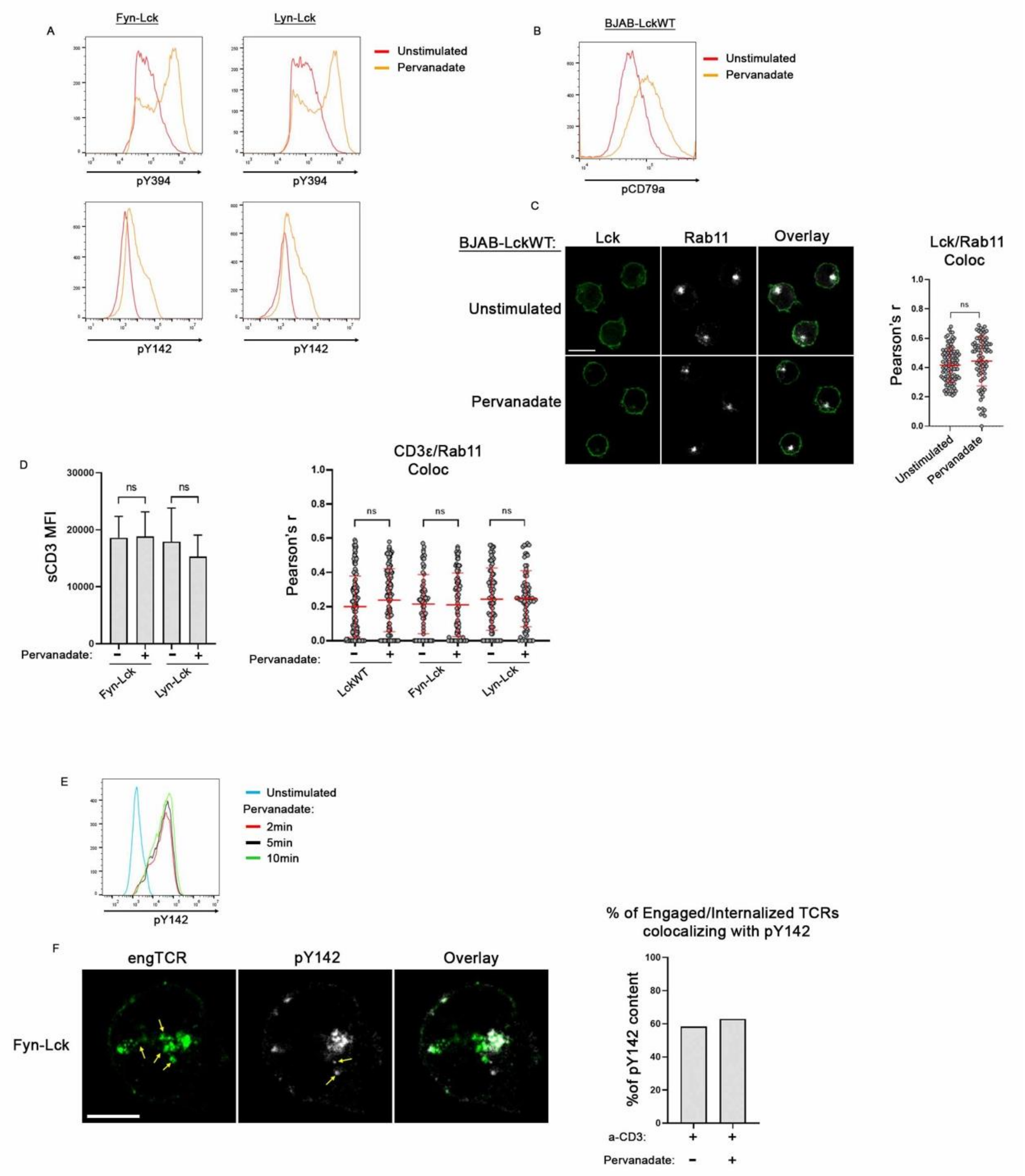


### Figure S3 (Corresponding to Fig.3)

A. Verification of Pervanadate-induced stimulation. Representative overlay histograms of Fyn-Lck and Lyn-Lck cells left untreated or incubated with 100μM Pervanadate for 10min and stained for pY394 and pY142 as indicated.

(B+C). Pervanadate treatment does not promote Lck translocation to the Rab11$^{+}$ EC in BJAB-LckWT cells. BJAB-LckWT cells left untreated or incubated with 100μM Pervanadate for 10min. B. Verification of Pervanadate-induced stimulation. Overlay histogram of the indicated samples treated as in (A) and stained for the phosphorylated ITAM Tyrosine, Y182, of the BCR CD79α (Igα) chain. C. Low magnification confocal images of cells stained for Lck and Rab11 and corresponding colocalization analysis graph. n≥70 cells from two independent experiments. Scale bar, 10μm.

D. Pervanadate stimulation does not promote TCR internalization or colocalization of CD3ε with Rab11.*Left panel*: Indicated cell lines treated as in (A) were stained for sCD3 and analysed by FACS, graph depicts MFI values from 3 independent experiments. *Right panel*: Collective colocalization analysis between CD3ε and Rab11 for the indicated cell lines treated as in (A). Images are not displayed as the staining patterns do not differ from images shown in Fig.2B

E. 2min of Pervanadate treatment suffice to reach saturation of ζ Y142 phosphorylation. Overlay histogram of Fyn-Lck expressing cells either left untreated or incubated with 100μM Pervanadate for the indicated time points, stained with a-pY142 and analysed by FACS.

F. Verification of ζ dissociation from a fraction of engaged TCRs after a-CD3 stimulation. Fyn-Lck cells were stimulated with a-CD3 for 10min or with a-CD3 + Pervanadate (100μM Pervanadate was added during the last two minutes of incubation to enable indirect detection of all ζ molecules within the cell) and stained with a-mouse $2^{ary}$ antibody (green) and pY142 (grey). *Left panels*: Representative high magnification confocal images. Yellow arrows, in corresponding staining panels, indicate either engaged/internalized TCRs lacking pY142 colocalization or $pY142^{+}$ spots without associated receptors, respectively. Scale bar,5μm. The proportions of engaged/internalized receptors containing pY142 fluorescence were quantified for each stimulation setting, to generate the adjacent graph. For both conditions approximately 40% of engaged/internalized TCRs were uncoupled from the ζ chain, in agreement with previous reports (15).

Numbers of engaged/internalised TCRs analysed: 264 (a-CD3 stimulation) and 156 (a-CD3+Pervanadate), pooled from three independent experiments

Unpaired Student t test; mean +/- SD; ns, not significant
